# Evaluation of the vectorial competence of *Glossina brevipalpis* in the transmission of *Trypanosoma congolense-*savannah type, in the Matutuíne District, Maputo Province, Mozambique

**DOI:** 10.1101/2025.07.24.666565

**Authors:** Nióbio V. Cossa, Fernando C. Mulandane, Denise R.A. Brito, Hermogénes Mucache, Moeti O. Taioe, Alain Boulangé, Geoffrey Gimonneau, Johan Esterhuizen, Marc Desquesnes, Luís C. B. Neves

**Affiliations:** Centro de Biotecnologia, Universidade Eduardo Mondlane, Maputo, Mozambique; Faculdade de Veterinária, Universidade Eduardo Mondlane, Maputo, Moçambique; Epidemiology, Parasites & Vectors, Agricultural Research Council-Onderstepoort Veterinary Research (ARC-OVR), Onderstepoort, South Africa; Unit for Environmental Sciences and Management, North-West University, Potchefstroom, South Africa; Intertryp, IRD, CIRAD, University of Montpellier, F-34398 Montpellier, France; Intertryp, CIRAD, Bouaké, Côte d’Ivoire; UMR Intertryp, CIRAD, Dakar, Senegal; Insitut Sénégalais de Recherches Agricoles, Laboratoire National de l’Elevage et de Recherches Vétérinaires, route du Front de Terre, 11500 Dakar-Hann, Sénégal; National Veterinary School of Toulouse (ENVT), 23 chemin des Capelles, 31076 Toulouse, France; Department of Veterinary Tropical Diseases, Faculty of Veterinary Science, University of Pretoria, Onderstepoort, South Africa

**Keywords:** Bovine, Buffy coat technique (BCT), Polymerase chain reaction (PCR), Cyclical transmission, Dissection

## Abstract

African Animal Trypanosomosis (AAT) is a parasitic disease caused by trypanosome species, posing a major challenge to animal health and production in Africa. In the Matutuíne District, southern Mozambique, where *Glossina brevipalpis* is prevalent, AAT is endemic. Previous research conducted in South Africa suggested that G*. brevipalpis* plays a marginal role in the transmission of *Trypanosoma congolense*. However, considering the wide distribution of *G. brevipalpis* in Southeast Africa and the major epidemiological implications of the aforementioned observations, this study was designed to re-assess the vectorial competence of *G. brevipalpis* in the transmission of *T. congolense-*savannah type.

The research involved 915 *G. brevipalpis* flies, fed six times on two cattle infected with *T. congolense*. After 48 hours, the flies were later fed on a non-infected animal to clean the proboscis, followed by extended feeding on four susceptible cattle for 22 days. These five clean animals were monitored for 60 days based on body temperature, packed cell volume, buffy coat technique and polymerase chain reaction. In the meantime, flies were dissected weekly to observe the development of trypanosomes in the midgut and proboscis. To discard any possible mechanical transmission by *G. brevipalpis* 48 hours after an infected blood meal, 97 flies from a colony were fed once on two infected cattle and once on a susceptible bovine. The Spearman correlation and Mann-Whitney U tests were used to analyse the data.

Results showed that the four susceptible cattle became infected with *T. congolense*, while the animal used for proboscis cleaning and discard any mechanical transmission remained uninfected. Dissections revealed that 89% of flies were positive for trypanosome through microscopy and 100% through PCR. These findings confirm that *G. brevipalpis* is a competent biological vector for transmitting *T. congolense,* challenging earlier assumptions and emphasizing its role in AAT dynamics.

## Introduction

African Animal Trypanosomosis (AAT), a parasitic disease caused by protozoa of the genus *Trypanosoma*, constitutes one of the main constraints on animal health and production, affecting several livestock and other domestic animals [1–4]. The consequent drastic reduction in productivity has a negative impact on farmers’ incomes (mainly small-scale farmers), who depend on this activity for their subsistence [5].

In sub-Saharan Africa, AAT in domestic animals is associated with four main etiological agents: *Trypanosoma congolense* (savannah, forest and Kilifi), *T. vivax, T. brucei brucei* and *T. simiae* [6–8]. Among these, *T. congolense*-savannah type is the species with the greatest veterinary importance, considering its wide distribution and its marked pathogenic effects on cattle [9–12].

The epidemiology of AAT is largely determined by the distribution and vectorial competence of its cyclical vector, the tsetse fly (*Glossina* spp.) [13–15], which infests an area of approximately 9 million km^2^ in Sub-Saharan Africa [1,16–18]. There is little information about the origin of the tsetse fly and trypanosomes complex, but it is believed that trypanosomes were probably intestinal commensals of primitive insects approximately 35 million years ago [19]. The first evidence of this complex was reported in 1903 and described by Bruce through the demonstration of the complete cycle of biological development of *T. brucei* within the tsetse fly [20]. These complex interactions allowed for adaptation, cyclical development of trypanosomes in the fly and a differentiated vectorial competence among tsetse fly species [13,21,22]. Differences in vectorial competence between *Glossina* spp. are influenced by the presence in the midgut, of different bacterial symbionts acquired from the environment [23–25]. This symbiosis ensures the survival and morphological development of the trypanosome through the synthesis of N-acetyl glucosamine, which inhibits a lectin lethal to the pro-cyclic forms of the parasite [26–29]. Among the three major microorganisms (*Wigglesworthia glossinidia*, *Sodalis glossinidius* and *Wolbachia pipientis*) [30], *S. glossinidius* is considered one of the most prominent due to its notable specific interaction with *Glossina* spp. [31–33], as well as its vertical transmission to larvae, during intra-uterine development [31,34,35].

The vectorial competence of *Glossina* spp., has already been studied in some parts of the African continent; in the case of *Glossina pallidipes* and *G. morsitans centralis*, they are considered competent vectors for the transmission of *T. vivax* [36]. Additionally, *G. m. centralis* is known to be a competent vector in the transmission of *T. congolense* and *T. b. brucei* [36,37]. In Mozambique, the first attempt to evaluate the vectorial competence of *G. brevipalpis* and *G. morsitans* in the transmission of *T. congolense* was carried out in 1947 by Hornby. In this experiment, there was a failure in breeding *G. morsitans* in captivity and inconclusive results regarding the vectorial competence of *G. brevipalpis* [38].

A study published by Motloang et al., [39] on the vectors of trypanosomes in KwaZulu-Natal strongly suggests a marginal role, if any, for *G. brevipalpis* in the transmission of *T. congolense*. Moreover, this paper attributes to *G. austeni* the exclusive role of transmission in this tsetse belt. It is worth mentioning that this study was conducted using laboratory-reared flies (not from South African origin) and *T. congolense* isolates from KwaZulu-Natal. Considering the large belt infested by *G. brevipalpis* from East Africa to North-East KwaZulu-Natal in South Africa and the major epidemiological implications of the aforementioned observations, this study was designed to reassess the vectorial competence of *G. brevipalpis* in the transmission of *T. congolense-*savannah type, using, for this purpose, wild flies from Matutuíne district, Mozambique and a *T. congolense*-savannah type isolated from the same geographic area and characterized with the 18S PCR-RFLP technique, for small subunit of ribosomal DNA.

## Methods

### Study site

The present study was conducted at the Tintigala Research Station (TRS) and the Biotechnology Center - Eduardo Mondlane University (CB-UEM) laboratory. The station is located in the district of Matutuíne, Zitundo Administrative Post, approximately 5 km from the southern gate (Gala gate) of the Maputo National Park (MNP), with geographic coordinates “-26.646158 S and 32.840419 E”. Matutuíne district is in the province of Maputo (south of Mozambique) between parallels 26° and 27°S and meridians 32° and 33° E.

### Fly-proof stable

A fly-proof stable with a total area of 39.1 m^2^ was built at TRS. The stable was divided into two compartments, one with an area of 21.5 m^2^ dedicated to the accommodation and feeding of cattle, and the remaining 17.6 m^2^ for experimental and therapeutic interventions.

### Experimental animals

Eight mixed (Nguni x Brahman) cattle, aged between 1 and 2 years, were brought from an area with no record of Animal Trypanosomosis (AT) occurrence. The farm of origin is located in the Province of Maputo, with the geographic coordinates S-25.92370 and E-E-32.51461. Before their inclusion in the experiment, all animals had their packed cell volume (PCV) checked and were tested against AT, using the buffy coat technique (BCT) [40] and polymerase chain reaction (PCR) [41]. The animals were then transported to the TRS in a fly-proof truck to prevent possible infections during the transport. Table 1 shows the role each animal allocated to the experiment.

**Table 1:** Different roles for the experimental animals. Legend. : Bovine **1** and **2** = experimentally infected by needle injection; **3** = used for proboscis cleaning; **4**, **5, 6**, **7** = used for the feeding of putatively infected flies and **8** = used for discarding any possible mechanical transmission experiment.

| Bovine | Experiment | Role |
| --- | --- | --- |
| **1** | Vectorial competence of *Glossina brevipalpis* | Infected donors |
| **2** |  |  |
| **3** |  | Non-infected animal used for proboscis cleaning |
| **4** |  | Clean receivers |
| **5** |  |  |
| **6** |  |  |
| **7** |  |  |
| **8** | Discard any possible mechanical transmission of *T. congolense* | Clean receivers |

### Experimental infection of cattle

For the experimental infection of cattle, a *T. congolense-*savannah type isolate (Isolate *CB* Matutuíne*-Gala* TCM 2018) from the CB-UEM cryobank was used. This isolate was collected in the study area in 2018 and cryopreserved by slow freezing in liquid Nitrogen (N_2_), using glycerol as a cryoprotectant. Two experimental animals (donors: bovine 1 and 2) were randomly selected and infected with 2 ml of infected blood diluted in glucose-phosphate saline solution (PSG), with a concentration of 10^4^ trypanosomes/ml. The animals were infected intravenously using a syringe with a volume of 2.5 ml and a 21G x 1/2^"^ gauge needle. The animals were monitored daily using BCT and PCR. Packed cell volume (PCV) was also monitored daily.

For the amplification of the trypanosome DNA in the blood sample of cattle, PCR targeting the ITS-1, a region of ribosomal DNA (rDNA), was conducted on the thermocycler T100 (BIORAD, USA) following the protocol from Njiru et al., [41], under the cycling conditions described in Table 2. For this, the reaction was carried out in a final volume of 20 μl, containing 1X Phusion Flash High-Fidelity PCR Master Mix (Thermo Fischer Scientific, USA), 0.5 *p*mol of each primer, 2.5 μl of eluted DNA (template DNA concentration of 13,75ng/μl-53,75 ng/μl) and distilled water was added to obtain a final volume of 20 µl. The primers used were ITS1 CF (5^’^-CCGGAAAAGTTCACCGATATTG-3^’^) and BR (5′-TTGCTGCGTTCTTCAACGAA A-3′) [41].

**Table 2:**
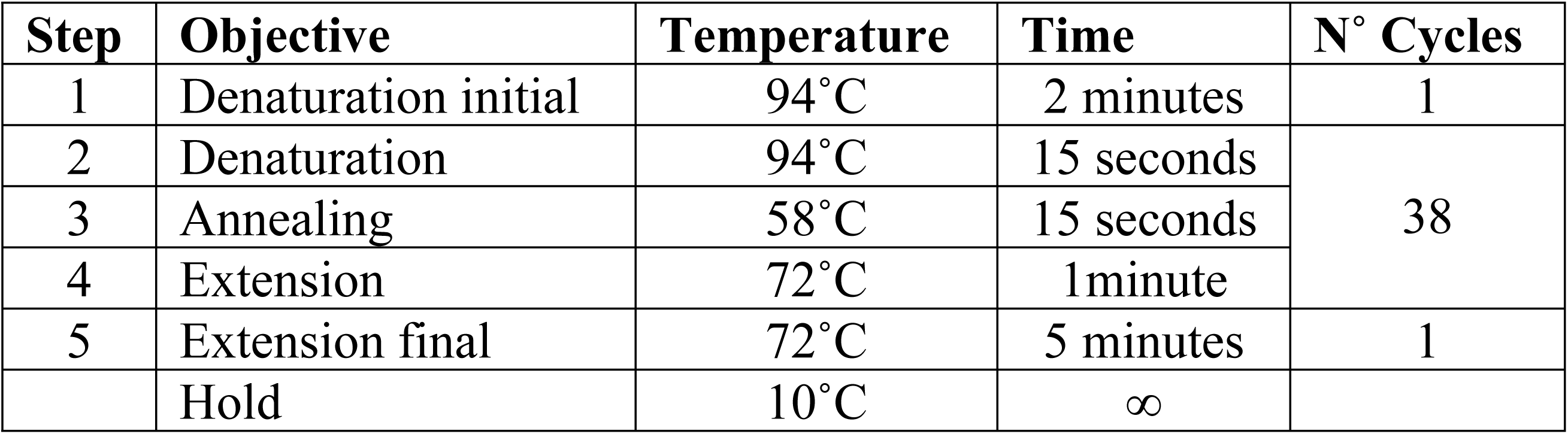
PCR conditions for amplification of the ITS1, a region of rDNA.

| Step | Objective | Temperature | Time | N° Cycles |
| --- | --- | --- | --- | --- |
| 1 | Denaturation initial | 94°C | 2 minutes | 1 |
| 2 | Denaturation | 94°C | 15 seconds | 38 |
| 3 | Annealing | 58°C | 15 seconds |  |
| 4 | Extension | 72°C | 1minute |  |
| 5 | Extension final | 72°C | 5 minutes | 1 |
|  | Hold | 10°C | ∞ |  |

### Fly capturing

Two days after the detection of infection in the two donor cattle, with motile *T. congolense* observed in the blood of the experimentally infected animals, a total of 915 *G. brevipalpis* flies (41 females and 874 males) were captured over a period of four days in areas close to the south entrance (Gala gate) of the MNP. Captures were conducted twice a day (morning and afternoon) using entomological nets (during car fly rounds) and H-traps [42]. Six traps, baited with 1-octen-3-ol and acetone, were deployed in three trapping points, separated by distances of at least 250 m [43,44]. Figure 1 illustrates the trap location and the net capture points, georeferenced using a Garmin GPS (GPSMAP®76). After capture and visual identification, the flies were placed in 2 types of cylindrical entomological cages (type 1: 19 cm diameter and 5 cm height; type 2: 10 cm diameter and 5 cm height) and transported to the TRS. The captured flies were grouped by date of capture and divided into groups of 10 or 25 flies, according to the cage size. To evaluate the vectorial capacity of *G. brevipalpis,* the flies were first starved for 48 hours and then allowed to feed on donor cattle every 48 hours, from day 4 to day 16 of patent infection, resulting in a total of 6 feedings. For this, the cages were placed against the animal’s body (two at a time against the dorsal and abdominal regions) for 5 minutes to allow the flies to feed.

**Figure 1:**
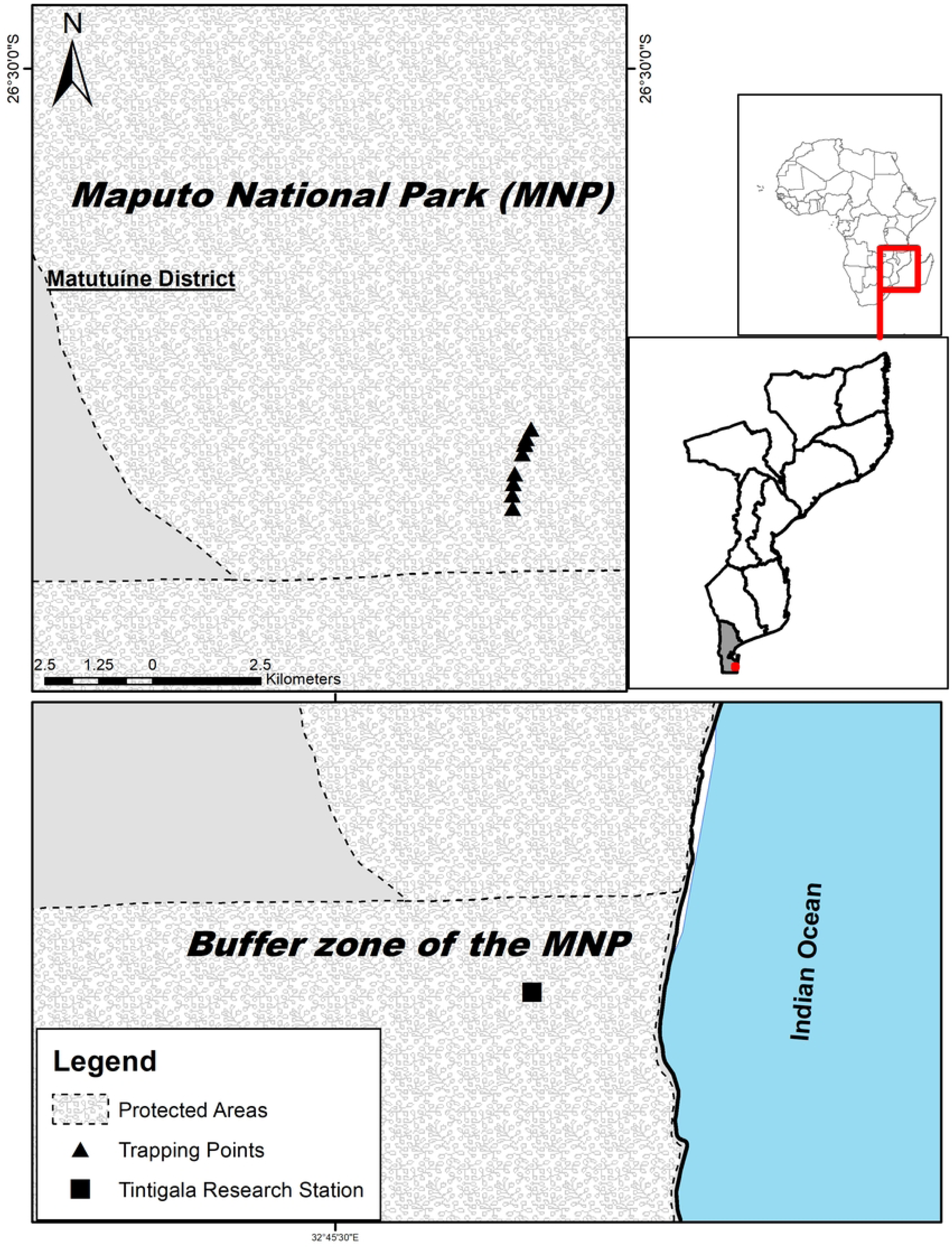
**Georeferenced fly capture points using H-traps and Entomological nets and Tintigala Research Station in Matutuíne District, Maputo province, Mozambique.**

### Cattle infection

Two days after the last feeding on donor animals, the flies were fed twice on one non-infected animal (proboscis cleaning: bovine 3) with a 48-hour interval between feeds, to remove any possible remnant trypanosomes in the proboscis from previously infected blood meals. Then, flies were fed each two days on the four clean receivers (susceptible animals: bovine 4, 5, 6 and 7), until motile trypanosomes were observed by BCT in their blood, and flies were then euthanized.

The five animals (the non-infected animal and the four receiver animals) were then monitored for 60 days, starting from four days after the last meal. During this period, the development of parasitaemia was monitored through clinical signs (fever, mucosal discolouration, lymph node inflammation, and weight loss), and laboratory tests using BCT, PCR, and PCV. For welfare reasons, animals diagnosed as anaemic, with PCV values below 18% for two consecutive days, were treated with diminazene aceturate at 7 mg/kg. At the end of the experiment, all animals were treated and later euthanized to avoid the accidental release of trypanosome isolates.

### Detection of infection in tsetse flies

To check whether tsetse flies are infected after the first blood meals on infected animals and observe the morphological evolution of *T. congolense* in the flies, a phased dissection of the flies was carried out to visualize the trypanosomes using the method described by Lloyd and Johnson (1924) quoted by Leak et al. [45]. For this, flies from 1 or 2 cages (20 flies) were dissected weekly, starting with the midgut on day 14 after the first meal, followed by the proboscis on days 21, 28 and 35.

To observe the trypanosomes, the proboscis and the midguts were placed on microscope slides containing droplets of PSG and covered with a coverslip. These organs were observed directly using a light microscope (10x eyepiece and 25x objective) and were then transferred to 1.5 ml tubes containing 70% ethanol and stored in a -20°C freezer for molecular analyses. For this purpose, DNA was extracted using the conventional precipitation method - ammonium acetate [46]. For the amplification of the trypanosome DNA in the flies, semi-nested 18S PCR was run, targeting a fragment of the 18S Ssu-rDNA as described by Geysen et al.[47], on a thermocycler T100 (BIORAD, USA) to supplement ITS-1 PCRs [47,48]. The first-round PCR was done using the forward primer 18ST nF2 (CAA CGA TGA CAC CCA TGA ATT GGG GA) and 18ST nR3 (TGC GCG ACC AAT AAT TGC AAT AC) as reverse primer. A second-round semi-nested was done using forward primer 18ST nF2 of the first amplification with the reverse primer 18ST nR2 (GTG TCT TGT TCT CAC TGA CAT TGT AGT G) [47].

### Discard any possible mechanical transmission of *Trypanosoma congolense* by *Glossina brevipalpis* after 48 hours from the last infected blood meal

While performing the previous experiments, 146 pupae of *G. brevipalpis* were obtained from 200 females (not included in the previous experiments) and placed in entomological cages for emergence. A total of 92 viable flies emerged in three batches of 37, 33 and 22 flies, producing trypanosome-free *G. brevipalpis.* Three days after emergence, each batch was fed once on one of the donor animals (i.e. animals infected during the previous experiment; bovine 5 and bovine 7). Forty-eight hours later, they fed once on a susceptible animal (bovine 8; clean receiver). This animal was then monitored for 60 days, starting four days after the last meal of the last batch.

### Statistical analysis

The data was collected and organised in Excel (Microsoft, USA) and all statistical analyses were done using STATISTICA v.13.3 (TIBCO Software Inc. 2017). The data recorded during the experiment did not present a normal distribution, so non-parametric tests were used for statistical analysis. A Spearman correlation test was used to analyse the variables body temperature, PCV and parasitaemia, with a significance level of 0.5% (p < 0.05). A comparative analysis of the prevalence of infection between the proboscis and midgut was carried out, using the Mann-Whitney U test. Finally, the Mann-Whitney U test was used to compare Microscopy and PCR results.

## Ethical considerations

Ethical approval for the experiments was obtained from the Research Ethics Committee of the University of Pretoria, Faculty of Veterinary Science (Ethics reference N°. REC 113-24) and Animal Research Ethics Committee of the Biotechnology Center-Eduardo Mondlane University (Ethics reference CEPA-CBUEM 07/2023).

## Results

A total of 915 *G. brevipalpis* were captured over 4 days (41 females and 874 males); most flies (95%) were captured using the fly-round technique (866 flies, 100% males). Using the stationary H traps, 49 flies (5%) were captured, out of which 84% were females.

### Experimental infection of cattle

After the infection of the two donor cattle, trypanosomes were detected by PCR on the 3^rd^ day post-infection (Figure 2). On the 5^th^ day, motile trypanosomes were observed in their blood, using BCT, showing a parasitaemia of 2 to 4 trypanosomes per preparation.

**Figure 2:**
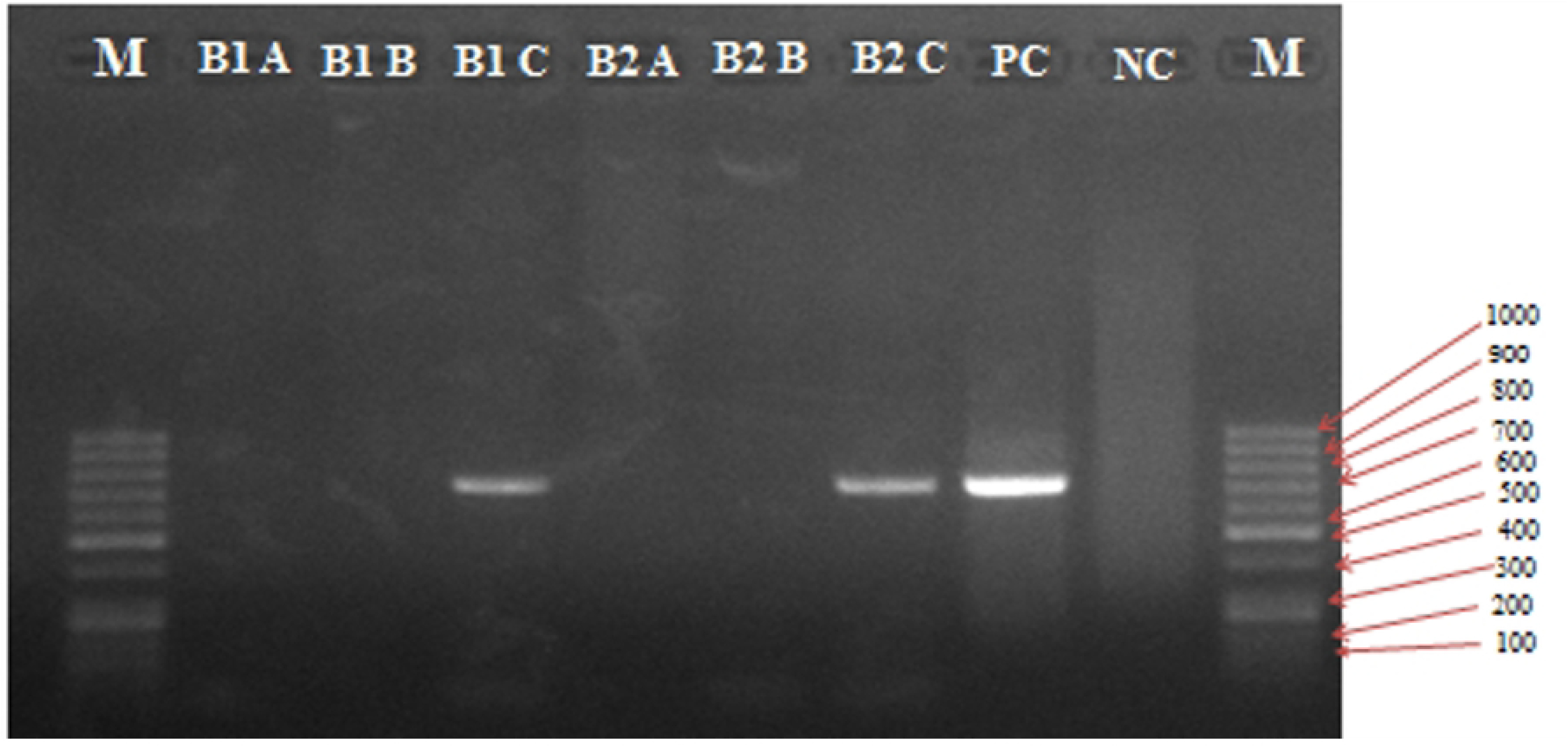
PCR product resulting from amplifying the ITS1 region of *Trypanosoma congolense*-savannah visualised in the 1.5% agarose gel, from the 2 donor cattle during post-experimental infection monitoring. Legend: M-100bp ladder; PC-Positive control (*T. congolense*-savannah isolated in blood of cattle); NC-Negative control; B1-Bovine 1; B2-Bovine 2; A-One day after experimental infection of donor cattle; B-Two days after experimental infection of donor cattle; C-Three days after experimental infection of donor cattle.

### Infection in susceptible animals (cleaning and receiver animals)

#### Non-infected anima (proboscis cleaning)

Captured flies were initially fed six times (over a period of 12 days) on infected cattle (donors) and subsequently fed twice (over a period of 3 days) on a non-infected animal (bovine 3). This animal (bovine 3) was monitored for 60 days, during which a body temperature variation of 38.1°C - 39.4°C was recorded. As for the PCV, it gradually increased, from 26% to 30% and no trypanosomes were observed using BCT or PCR, as illustrated in Figure 3.

**Figure 3:**
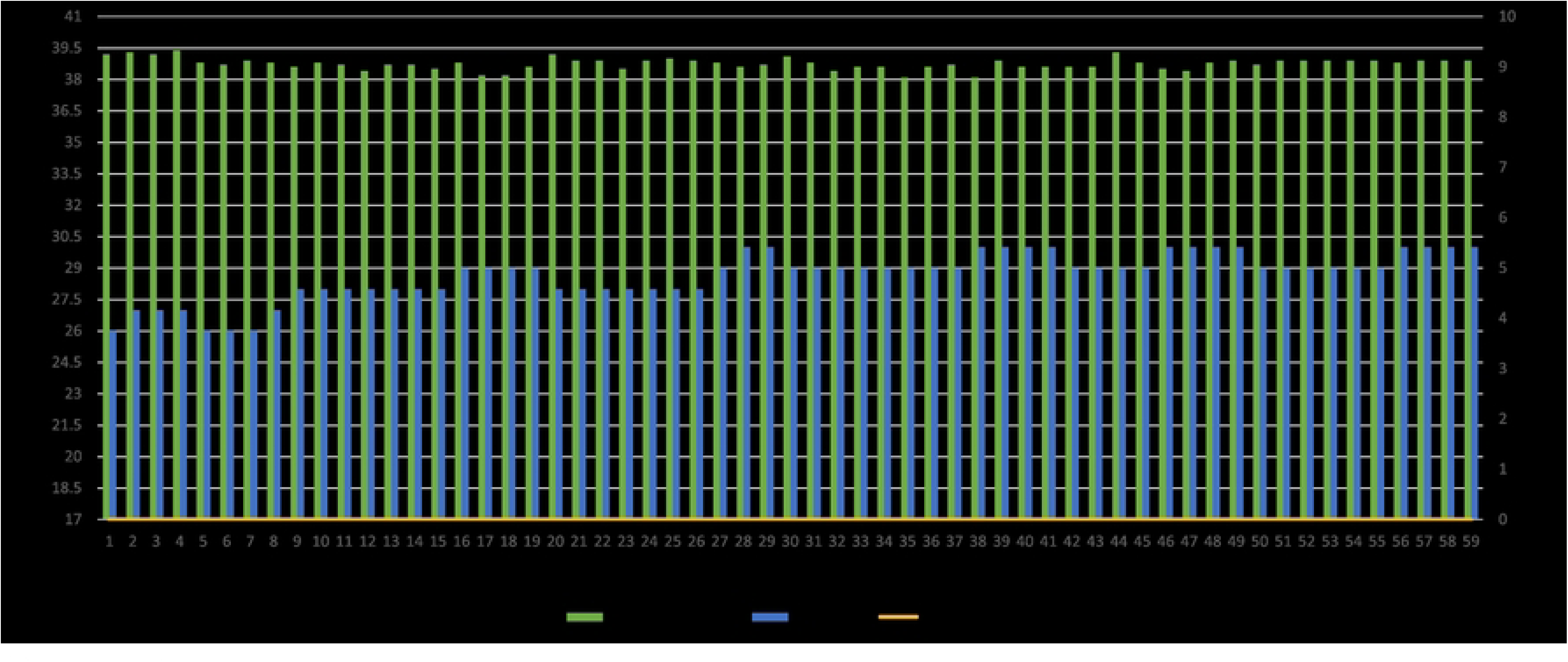
**Record of temperature, PCV and parasitaemia for the cleaning animal (Bovine 3).**

#### Receiver animals

Monitoring of all four receiver animals (bovine 4, 5, 6 and 7) started on the 4^th^ day after the first meal by infected flies. Subsequently, all receiver animals became infected with an incubation period varying from 8 to 22 days, presenting considerable changes in the three main monitored parameters (average body temperature, PCV and parasitaemia), as illustrated in Figure 4. The monitoring period for the development of parasitaemia ranged from 17 to 40 days and was determined by the PCV level (≤ 18%) recorded in each animal.

**Figure 4:**
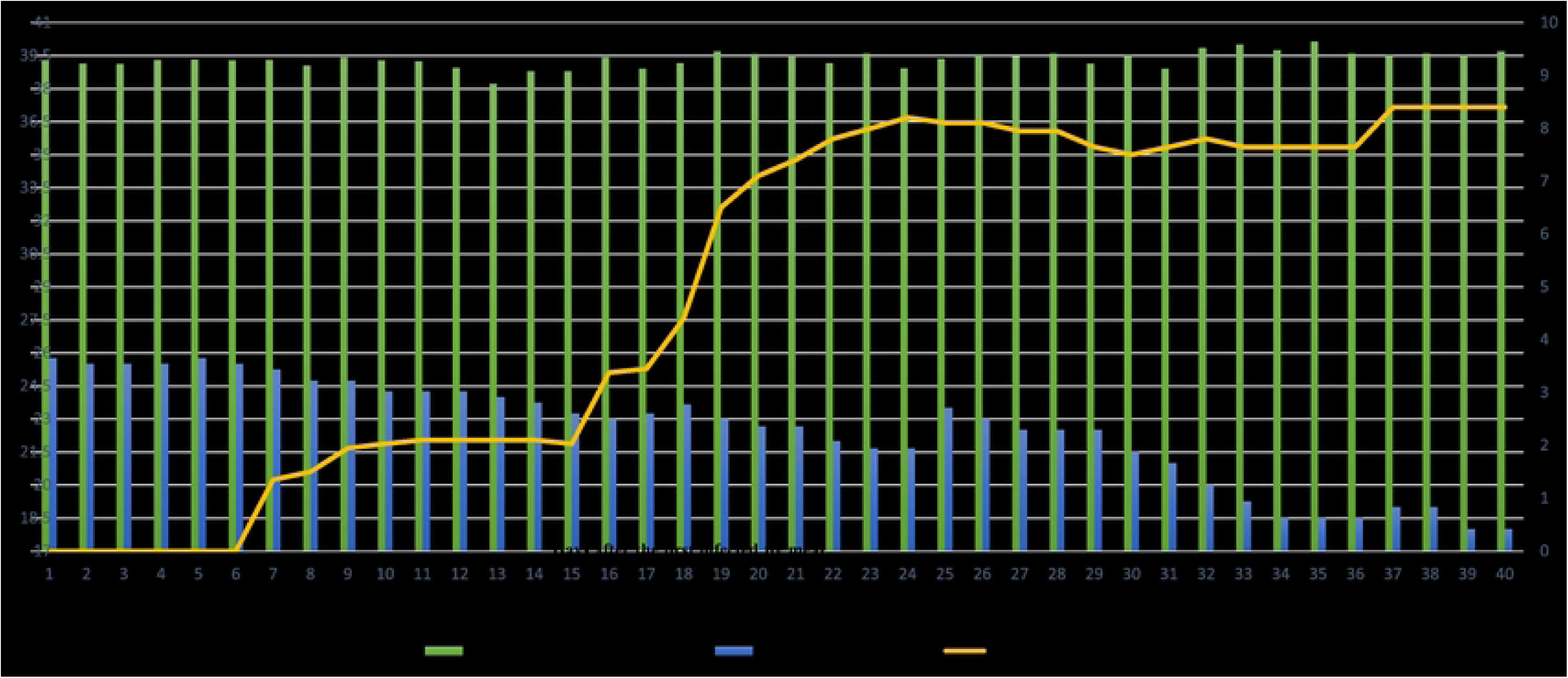
**Records of the average temperature, PCV and parasitaemia for receiver animals.**

The prepatent period in the four receiver animals differed, varying from 10 to 22 days. The observed level of parasitaemia ranged from 2 to 7 trypanosomes/preparation and a maximum peak of parasitaemia estimated at 8.4 on the Herbert and Lumsden scale was recorded. The infestation was also characterised by intervals of high (8.4) and low (7.5) parasitaemia.

The PCV of the four receiver animals was characterised by a stabilization period from days 4 to 11, followed by a gradual reduction on day 12 after the first infected fly meal until day 29, where a slight increase was recorded, followed by a gradual reduction again. A remarkable difference was observed concerning the beginning of the reduction in PCV levels, which was observed starting between days 9 and 19.

The body temperature of the same receiver animals was characterised by intermittent fever that ranged between 39.6°C and 40.9°C, with differences between animals. An average of four fever episodes was observed, the first (39.6°C) on day 23 and the last (39.7°C) on day 44 after the first infected fly meal.

The statistical analysis of the parameters recorded in the susceptible animals revealed a significant moderate negative correlation between parasitaemia and PCV (r = - 0.691358, p=0.001). Additionally, a significant positive correlation was observed between body temperature and parasitaemia (r = 0.436422, p=0.002).

### Discard any possible mechanical transmission of *Trypanosoma congolense* by *Glossina brevipalpis* 48 hours after the last infected blood meal

In this experiment to discard possible mechanical transmission, 48 hours after the last infected blood meal, the bovine (bovine 8) was monitored for 60 days without the occurrence of fever (body temperature variation: 38°C - 39.5°C). No significant reduction in PCV and no motile trypanosomes were observed through BCT or detected through PCR (Figure 5).

**Figure 5:**
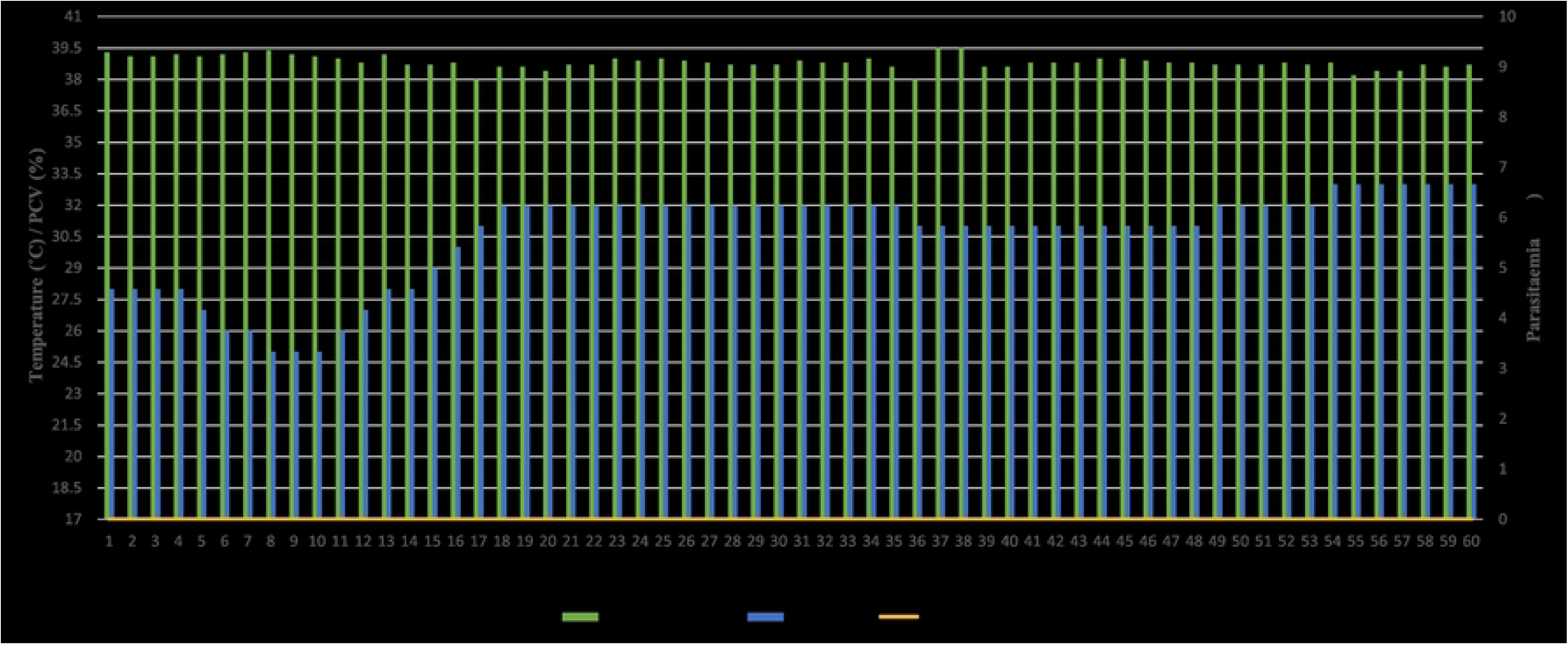
**Registered temperature, PCV and parasitaemia in bovine used for discard any possible mechanical transmission.**

It is worth mentioning that all samples from animals susceptible to *T. congolense* infection, whether collected for the analysis of vectorial competence (bovine 3, 4, 5, 6 and 7) or mechanical transmission (bovine 8), were also tested by PCR; the results are illustrated in Figure 6.

**Figure 6:**
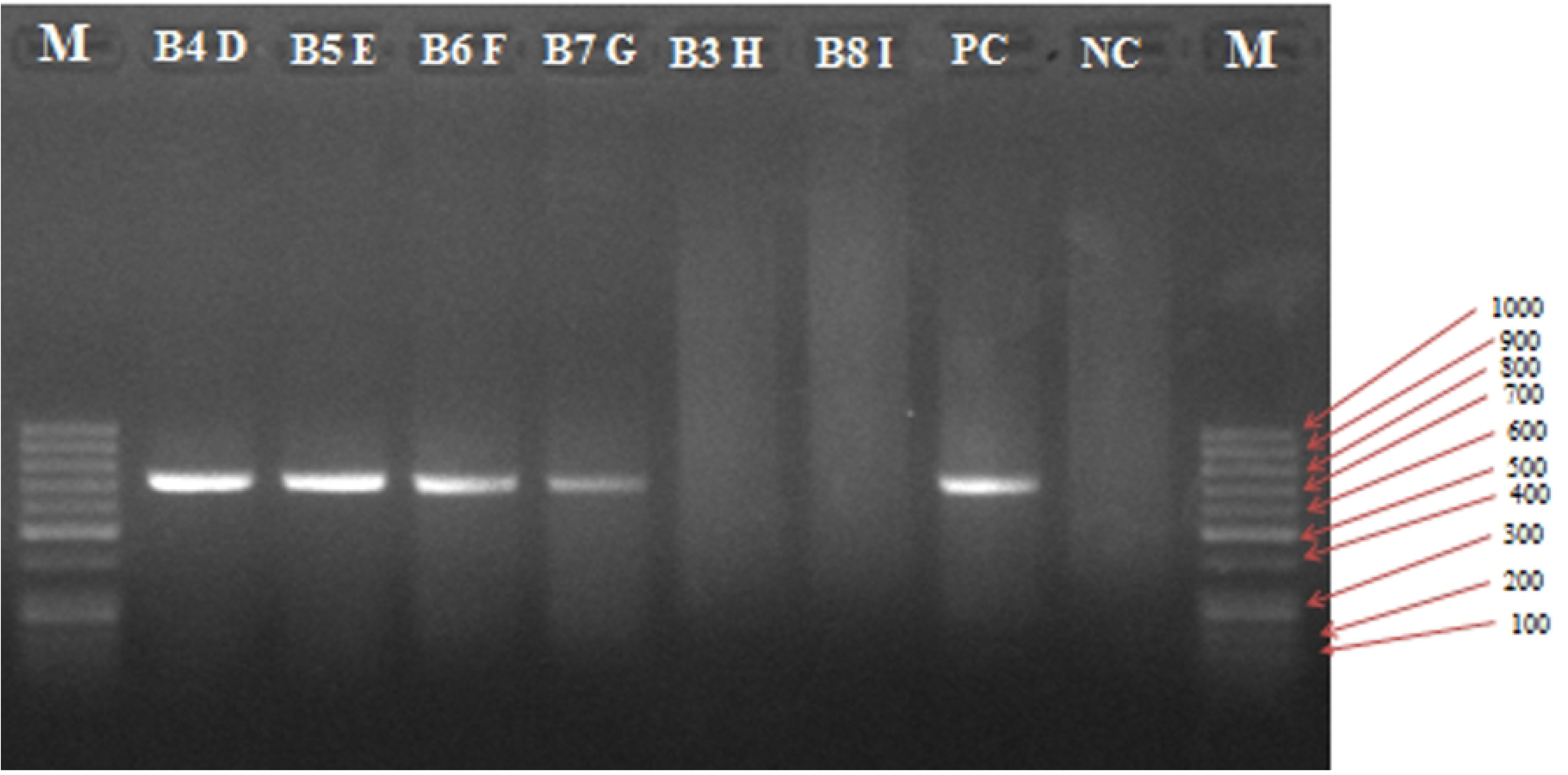
PCR product resulting from amplifying the ITS1 region of *Trypanosoma congolense*-savannah type visualised on a 1.5% agarose gel, from 6 susceptible cattle during post-experimental infection with flies. Legend: M-100bp ladder, PC-Positive control (*T. congolense*-savannah isolated in blood of cattle), NC-Negative control; 3,4,5,6,7 and 8-Identification of bovine, D-10 days after first infected fly meal in receiver animal, E-19 days after first infected fly meal in receiver animal, F-21 days after first infected fly meal in receiver animal, G-22 days after first infected fly meal in receiver animal, H-64 days after first infected fly meal in cleaning animal, 1-64 days after first infected fly meal in cleaning animal

### Determination of infection in tsetse flies

Using a light microscope, 71 flies (89%) were detected positive for the presence of motile trypanosomes and the remaining 9 flies (11%) tested negative. During dissection and microscopic observation of the midgut on day 14, the presence of motile trypanosomes was detected in 13 out of the 20 dissected flies (65%). On day 21, the dissection of the proboscis revealed 18 infected flies out of 20 dissected flies (90%). Finally, the proboscis was dissected at days 28 and 35, at which the presence of trypanosomes was observed in all 20 flies (100% infection) in both periods, as illustrated in Table 3.

**Table 3:** Microscopy and PCR results of dissected flies.

| Microscopy result |  |  |  |  | PCR result |  |  |
| --- | --- | --- | --- | --- | --- | --- | --- |
| Days | Organ | Positive (%) | Negative (%) | Total | Positive (%) | Negative (%) | Total |
| 14 | Midgut | 13/20 (65) | 7/20 (35) | 20 | 20/20 (100) | 0 | 20 |
| **21** | Proboscis | 18/20 (90) | 2/20 (10) | **20** | 20/20 (100) | 0 | **20** |
| **28** | Proboscis | 20/20 (100) | 0 | **20** | 20/20 (100) | 0 | **20** |
| **35** | Proboscis | 20/20 (100) | 0 | **20** | 20/20 (100) | 0 | **20** |
|  | **Total** | 71/80 (89) | 9/80 (11) | **80** | 80/80 (100) | 0 | **80** |

Molecular diagnosis (semi-nested 18S PCR) of the dissected flies revealed the presence of infection in the 80 dissected flies, tested from day 14 onwards (midgut) and proboscis at days 21, 28 and 35. Given its species-specific nature, this method allowed for the visualization of a band at about 600 bp and another at 750 bp in some samples, revealed in agarose gel in all samples tested (Figure 7). Statistical analysis revealed a significant difference (z = 2.48, p = 0.01) between the number of positives for trypanosomes detected using microscopy versus the number of positives detected using PCR.

**Figure 7:**
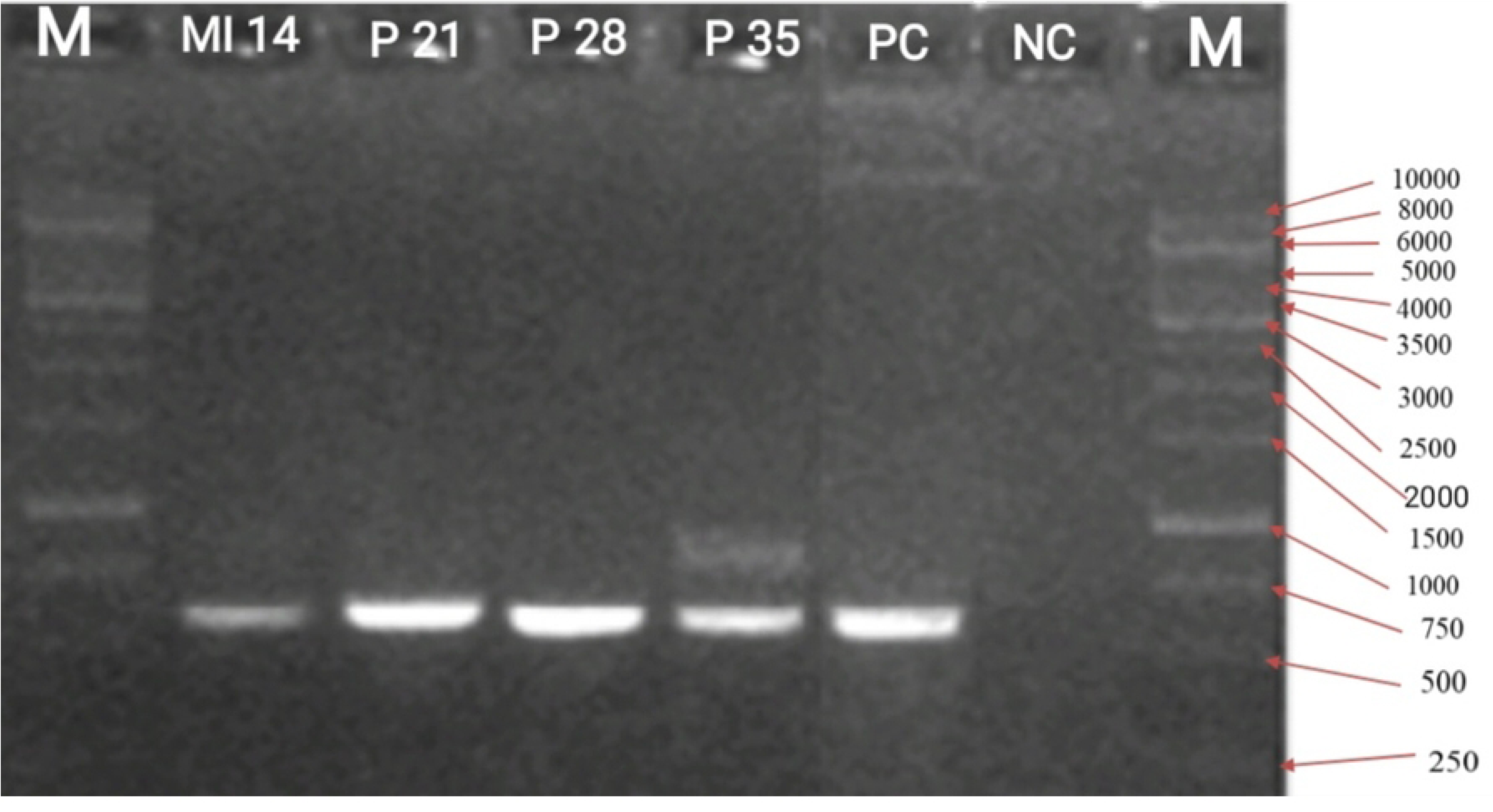
PCR product resulting from amplifying the 18S ribosomal DNA region of *Trypanosoma congolense* visualised on a 1.5% agarose gel, from 4 samples of *G. brevipalpis* after dissection. Legend: M-1kb ladder; PC-Positive control (*T. congolense* savannah isolated in blood of mice); NC-Negative control; MI = midgut ; P = proboscis; 14= samples of tsetse fly at day 14; 21= samples of tsetse fly at day 21; 28= samples of tsetse fly at day 28; 35 = samples of tsetse fly at day 35.

## Discussion

The complete elucidation of vector competence status in the transmission of pathogens is an essential component to unravel the epidemiological dynamics of vector-borne diseases. In this study, the ability of *Glossina brevipalpis* to transmit *Trypanosoma congolense*, which was questioned by Motloang *et al*., 2012 (there should be a reference here in hooks), was investigated.

A total of 915 wild flies were captured, using two different methods, the H trap and car fly round. One of the biggest disadvantages of using H traps to capture live flies for breeding purposes is the poor survival of captured flies. This may be associated with the exposure of flies to unfavourable temperatures and relative humidity inside plastic bottles, especially when flies remain inside for a long time. In order to improve fly survival, the use of mosquito net cages could be considered as an alternative to plastic bottles. The observed predatory behaviour of ants and spiders towards the trapped flies is another major drawback of this method.

It is generally accepted that the pre-patent period in experimental laboratory infections with trypanosomes in mice is 3 to 10 days [49], in cattle is 6 to 12 days [27], depending on the species of trypanosome involved, the immune status of the host and the infective dose [49, 50]. Studies carried out with *T. congolense* indicated a pre-patent period of 5 to 13 days [51–53]. These values are consistent with those observed in the experimental infection of donor cattle in this study, in which the parasites were detected by PCR on day 3 post-infection and using the BCT technique on day 5 post-infection (first observation of a motile *T. congolense* parasite). In addition to the high sensitivity of PCR, this early onset of parasitaemia is probably associated with the use of the intravenous route for the experimental infection [54], although some authors do not consider this as a determining factor for the length of the pre-patent period [51].

All four receiver animals, which were used to feed *G. brevipalpis* flies harbouring mature infections, showed patent *T. congolense* infections after an incubation period varying from 8 to 22 days. This observation is consistent with the interval of 4 days to 8 weeks, previously reported [55, 56]. One of the main clinical signs of animal trypanosomosis is the onset of fever, sometimes intermittent, associated with the onset and progression of parasitaemia in infected animals [27,57,58]. These clinical signs were clearly observed in the four susceptible cattle. The differences in time intervals at which the animals presented clinical signs were probably associated with their individual immune responses, given that they were all infected with the same trypanosome isolate in very similar doses. A progressive reduction in PCV was also recorded in the four receiver animals, confirming anaemia as one of the most typical symptoms of AT [39,59,60]. Anaemia is extensively reported in studies carried out with animals infected with *T. congolense* [27,52,61].

During the parasitaemia monitoring phase, fluctuations in this parameter and intermittent fever were observed, accompanied by very low levels and sometimes aparasitaemic episodes. This is a common phenomenon observed in the progression of this disease in ruminants [62,63], which is associated with the host immune system’s process of eliminating waves of new parasite populations emerging due to antigenic variation [64].

In an attempt to increase the time from the last infective blood meal and to avoid any possible mechanical transmission, the flies were fed twice on an un-infected to clean flies mouthpart, and this took place on day 11 and 13 after the flies had their first meal on the infected donor animals. At 11 to 13 days post-ingestion of *T. congolense* by the tsetse fly, the trypanosome is likely to be in the differentiation and multiplication phase at the level of the cibarium, which probably explains the absence of mature metacyclic trypomastigotes, which are necessary to initiate a host infection [65]. This fact justifies the non-detection of trypanosomes, or their classical clinical manifestations, during the 60-day monitoring period of the non-infected calf used for the cleaning of the proboscis, thus ruling out the possibility of a potential mechanical transmission or a regurgitation process at this stage of *G. brevipalpis* infection. Additionally, results of the mechanical transmission experiment using teneral flies indicate that *G. brevipalpis* cannot mechanically transmit *T. congolense* 48 hours after an infected blood meal. Nevertheless, a possible role of *G. brevipalpis* in mechanical transmission of *T. congolense* still need to be explored through interrupted feeding on infected cattle and restarting feeding on a potential receiver within 0-30 minutes, the period during which trypanosomes remaining on the mouthparts of biting flies are thought to be fully alive and infective [66,67].

In addition for the 4 receiver cattle, the dissection and microscopic observation of experimentally infected flies allowed for the detection of 89% of *G. brevipalpis* positive for trypanosome, while molecular diagnosis (PCR), using the 18S rDNA marker, allowed for the amplification of *T. congolense* DNA in 100 % of the flies tested according to the period of development and the dissected organ (either midgut or proboscis). These results confirmed the high sensitivity and specificity of PCR in the detection of trypanosomes in tsetse flies when compared to dissection. Results with significant differences between the two techniques were also reported by Ouma et al., [68], who reported a *T. congolense* infection rate of approximately 100% using PCR and an infection rate of <4% using dissection.

In addition to the demonstration of the vectorial competence of *G. brevipalpis*, recent studies have reported *T*. *congolense* infection in flies from the field with rates of 3.7% and 89% reported in populations from East and Southern Africa, respectively [69,70]. Furthermore, infection rates of 12.74 % and 3.19 % were recently reported in *G. brevipalpis* and *G. austeni,* respectively, in the Matutuíne district [71]. These studies demonstrate the occurrence of *T. congolense* infections in *G. brevipalpis.* However, they provide little information regarding the origin of the infection or the distinction between mature and immature infections. Studies investigating the maturation of *T. congolense* in wild *G. brevipalpis* reported a weak infection rate of 1.6% at the level of the proboscis [72]. Moreover, the occurrence of a patent infection was observed in rats after injection of the product resulting from the crushing of the proboscis detected positive to *T. congolense* [73].

In this study, the results obtained through phased dissection of *G. brevipalpis* and microscopic observation, thereafter confirmed by PCR, demonstrated a 100% infection rate in the midgut on day 14 and 100% in the proboscis on days 21, 28 and 35. This drastically contrasts with the results reported by Motloang et al., [39] in which there was only 2% infection in the midgut and no infection in the proboscis. The results recorded in the present study may be partly explained by 1/ the putative compatibility between the local trypanosome strain and the *G. brevipalpis* used in this experiment due the their similar geographic origin; 2/ the use of flies captured in the field (probably of different ages); 3/ the stress of hunger associated with the 48-hour interval between meals; 4/ the 10 days of patent infection observed before the first meal to allow the maturation of the trypanosome infection in donor animals; 5/ the number of meals (six) on donor cattle. Generally, the susceptibility of infection of tsetse flies by *T. congolense* is considered relatively high in adult flies compared to the teneral stage [74,75], and the ability to acquire infection is directly related to age, especially in females of the *G. brevipalpis* species [76]. Tsetse flies present greater transmissibility of the infection in strains isolated from domestic animals compared to those of wild origin [77]. It is worth mentioning that the *T. congolense* strain used in this study was isolated from domestic animals and passaged once in mice. Moreover, environmental stress and food deprivation have been reported as crucial factors contributing to the increased susceptibility of flies, whether young or adult, to infection by *T. congolense* and *T. brucei* [78–81] because of the reduction in the fly’s immune response to external agents, in particular, trypanosomes. Additionally, different species of wild *Glossina* spp. may harbour different genotypes of *Sodalis glossinidius* that may create a fertile environment for the biological development of trypanosomes [24]. However, its prevalence rate varies between tsetse fly species and geographic location [82,83]. A study carried out by Dieng et al., 2022 [82], in different African countries, including Mozambique, reported the occurrence of *S. glossinidius* at a rate of 14% in the wild population of *G. brevipalpis* from Matutuíne [82], though more in-depth studies on the prevalence and association with trypanosomes still need to be carried out.

## Conclusion

This study revealed a high infection of *G. brevipalpis* with *T. congolense*-savannah type at midgut and proboscis level, as well as the PCR-confirmed occurrence of *T. congolense*-savannah infections with their classical clinical signs in the four receiver cattle. These results demonstrated the ability of *G. brevipalpis* to acquire trypanosome infection, to effectively host the cyclical development of *T. congolense-*savannah, and to transmit the parasite to cattle. These findings reinstate *G. brevipalpis* as a competent vector of *T. congolense*-savannah and highlight its important epidemiological role in the maintenance of the *T. congolense* cycle in vast regions of South and East Africa.

## Acknowledgements

We would like to acknowledge the Biotechnology Centre of Eduardo Mondlane University for using their laboratory. We would also like to acknowledge Keila Zandamela, Edmilson Philimone, Simão Sitoe and Nelson for their assistance and dedication during this project. This project has received funding from the European Uniońs Horizon 2020 research and innovation programme under grant agreement n° 101000467, acronym "COMBAT” (Controlling and Progressively Minimizing the Burden of Animal Trypanosomosis).

## Authors’ contributions

All authors read, amended and approved the final manuscript. **Conceptualization**: Nióbio V. Cossa, Fernando C. Mulandane and Luís Neves. **Sampling**: Nióbio V. Cossa, Fernando C. Mulandane, Hermógenes N. Mucache. **Corrections**: Nióbio V. Cossa, Fernando C. Mulandane, Moeti O. Taioe, Alain Boulangé, Geoffrey Gimonneau, Johan Esterhuizen, Marc Desquesnes, Luís C.B. Neves. **Data analyses**: Nióbio V. Cossa and Denise R.A. Brito. **Maps and design of Figures**: Nióbio V. Cossa and Fernando C. Mulandane. **Writing of the original draft**: Nióbio V. Cossa, Fernando C. Mulandane and Luís Neves.

